# A survey of the Loureedia genus (Araneae, Eresidae) with a new species from Iran and the first assessment of illegal wildlife trade as a threat to the group

**DOI:** 10.1101/2020.05.07.082891

**Authors:** Sérgio Henriques

## Abstract

We describe and illustrate a new species of velvet spider from Iran, *L. venatica* sp. n., the first species of this genus outside of the Mediterranean region. We also resurrect *Eresus jerbae* from synonym, as a distinct and valid Loureedia species, *L.jerbae* comb.n, and record this genus to Jordan for the first time.

We map the distribution of all available observations of *L.venatica sp.n*. and *L.jerbae* comb.n, based on museum specimens and on photographic records, using these observations, and the uncertainty therein, to estimate the species range and how it would be classified under the IUCN Red List. Addressing two of the obstacles to the conservation of poorly known taxa, the Linnaean shortfall, by increasing the number of described species, and the Wallacean shortfall, by increasing current knowledge of species distribution as well as their range.

We also found that *Loureedia jerbae* comb.n. from Tunisia is been sold as an exotic pet, and that photos of Iranian *Loureedia venatica* sp.n. are being used to advertise the sale of this genus in the pet trade. We discuss the impacts this likely causes to these species, as well these species extinction risk under the IUCN Red List.

## Introduction

Eresidae, commonly known as velvet spiders, is a relatively small family consisted of only nine genera and 98 species known mostly from Africa and Eurasia, with a single representative from Brazil [WSC, 2020].

So far, three species of Eresidae belonging to *Eresus* and *Stegodyphus* genera have been recorded in Iran [Zamani et al., 2020]. The genus *Loureedia* although recorded to Iran (Henriques et al. 2018), has thus far not been taxonomically analyzed in the region and is the focus of the present work.

The Loureedia genus, currently contains three Mediterranean species spanning from Morocco To Israel, at a remarkable distance from Iranian records [Henriques et al. 2018]. We here present our analysis of Iranian Loureedia specimens for the first time and recognize that the male morphology (such as the fine structure of the male pedipalp and dorsal pattern) are not similar to any previously described species. While comparing Iranian species with other Loureedia we recognized that *Eresus jerbae* (El-Hennawy, 2005) solely known from females, is currently considered as junior synonym of *Loureedia annulipes* (Lucas, 1857) represents a separate species and therefore we resurrect it from synonymy.

*Lourredia jerbae* **comb. n**., **stat. resurr**. is morphologically the closest member of this genus to Iranian *Loureedia*. Revealing how important it is to have in depth knowledge of the group as a whole (*Loureedia* and *Eresus*), before describing new species, as the Iranian specimens which we were recognize as a news species (solely known from males) could have been the undescribed male of *L.jerbae*.

The goals of this paper is therefore to address the Linnaean shortfall the Wallacean shortfall (i.e. Bini et al. 2006; Brito, 2010) of this genus, by describing a new Loureedia species from Iran, resurrect Eresus jerbae as a valid species, validate it’s placement in the *Lourredia genus as L.jerbae*, provide comparative figures to allow the identification of these species and discuss their distribution, specifically in how it relates to IUCN Red List criteria as well relevant threats, particularly relating to their present in the pet trade.

## Methods

Morphological terminology follows Miller et al. (2012), and has been adapted to describe the fine morphological structures of the male pedipalp (Henriques et. al. 2018).

The holotype was photographed at the Zoological Museum of University of Turku, Finland, analysed using an Olympus SZX16 stereomicroscope coupled with an Olympus Camedia E-520 camera which assisted in preparing the illustrations. Digital images were stacked using Zerene Stacker version 1.04.

Lengths of leg segments were measured dorsally side and are presented as the total length (of femur, patella, tibia, metatarsus and tarsus. All measurements are given in millimetres.

Studied material is deposited at the Biologiska Museet, Lund, Sweden (BML), British Natural History Museum, London, UK (BNHM), Laboratory of Arachnid Cytogenetics at Charles University in Prague, Czech Republic (LACCUP), the Muséum d’histoire naturelle, Genève, Switzerland (MHNG), Muséum National d’Histoire Naturelle, Paris, France (MNHN),Natural History Museum of the University of Florence, Italy (UNIFI), Zoological Museum of University of Turku (ZMUT) and Sergio Henriques private collection (SHPC).

Abbreviations used for the eyes were ALE – anterior lateral eye, AME – anterior median eye, PLE – posterior lateral eye, PME – posterior median eye. Abbreviations used for the leg segments were: Fe – femur, Pt – patella, Ti – tibia, Mt – metatarsus, Ta – tarsus.

## Results

### Loureedia

Miller, Griswold, Scharff, Řezáč, Szűts & Marhabaie urn:lsid:zoobank.org:act:5FEC8D28-5F6F-4E58-A5C2-5EEBD35B0090 http://species-id.net/wiki/Loureedia

Type species. *Eresus annulipes* Lucas, 1857.

#### Current composition

Five species: *L. annulipes* (Lucas, 1857), *L. colleni* Henriques, Miñano & Pérez-Zarcos, 2018, *L. jerbae* (El-Hennawy, 2005), *L. lucasi* (Simon, 1873) and *L. venatica* **sp. n**.

*Loureedia spp*. occur from Morocco to the west and appears to have its eastern most reach in Iran (Henriques, 2018), which is remarkably Eastward from the nearest known location of the genus in Jordan. A new record to the country which we present here for the first time (Fig. 1).

**Figure 1.**
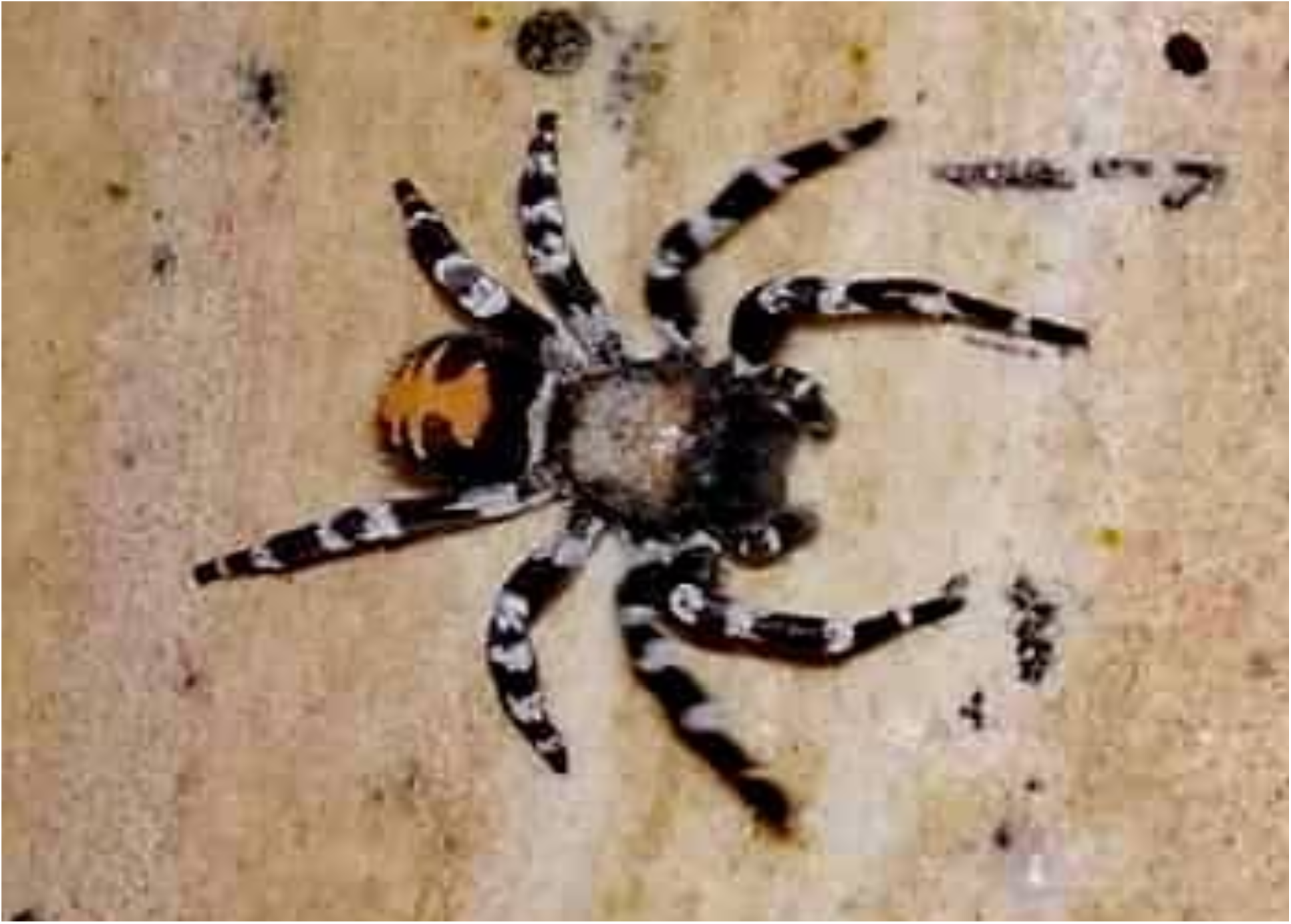
First *Loureedia* record from Jordan, Northern Azraq (photo by Majd al Qadi)

Although we were not able to analyse any Jordanian specimen at the time of this publication, the photographic record presented here can be confidently identified as a male Loureedia sp., as it presents the characteristic male coloration, not known in any other eresid genera (Miller et al., 2012; Henriques et al., 2018). The specimen was recorded on the October 30 (2018) in Northern Azraq, which matches the known phenology of other species within the genus (Henriques et al., 2018). The almost complete absence of white markings in dorsal pattern (Fig.1) could indicate that it might not belong to Loureedia annulipes (known from the neighbouring regions), and could potentially represent another undescribed species.

Given our present analysis of this genus, it would not be surprising to find this group in Iraq or neighbouring countries, but research efforts have been hampered by recent conflicts, and species studies and discovery has been all but halted in the region and are sadly not forthcoming. Conflict which is clearly in large part responsible for the Wallacean shortfall in the region

*Loureedia* have previously been recorded to Iran (Henriques et al. 2018: fig. 2h), but the taxonomy of the genus in the country has never been analysed; a Linnean shortfall which we will address.

**Fig. 2.**
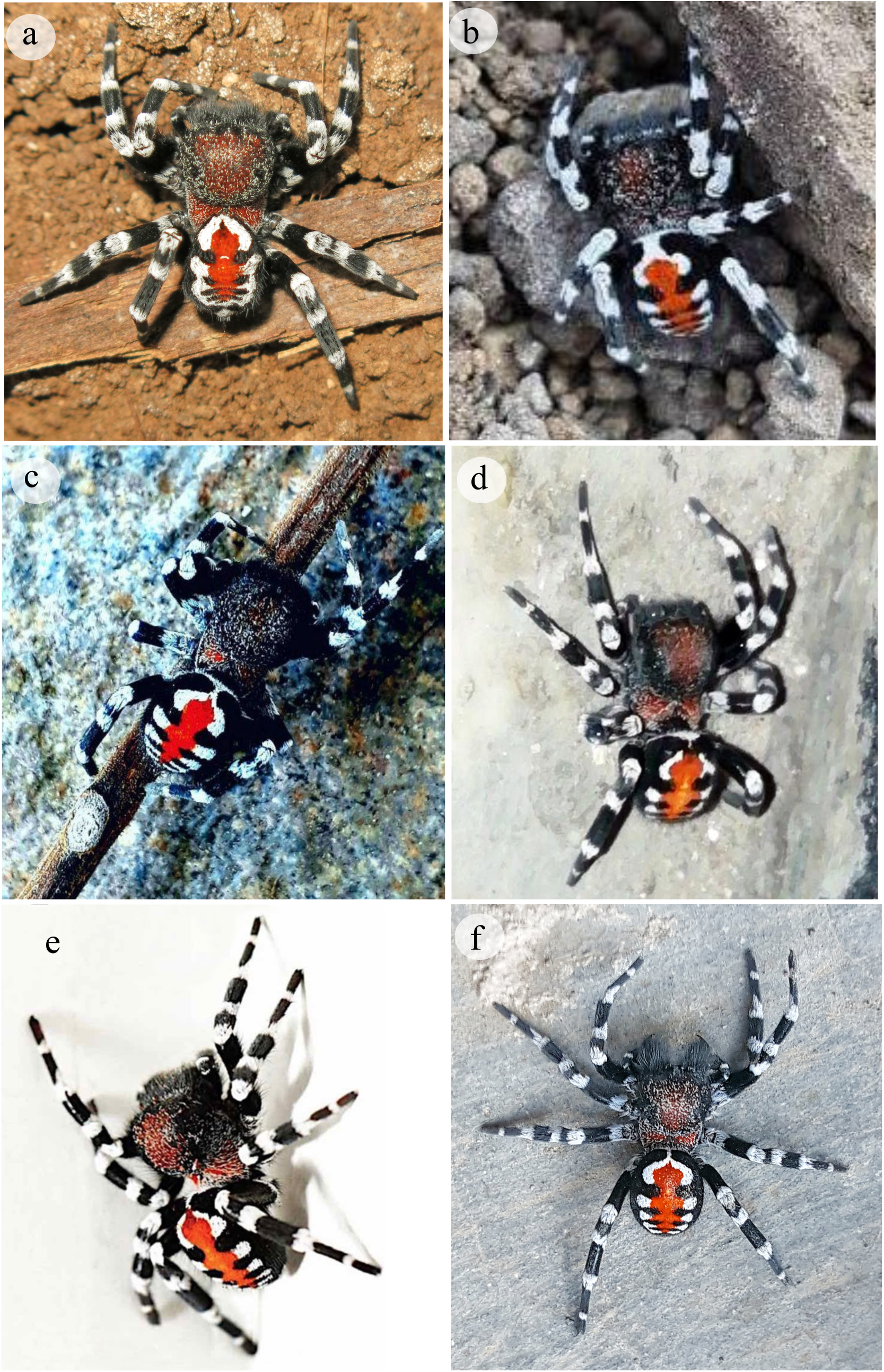
Photographic records of male *Loureedia venatica sp.n. in vivum* from the Iranian provinces of a) Tehran (photo by Alireza Zamani), b) Tehran (photo by Bayan Golavi), c) Kerman (photo by Hadis Asgari), d) Fars (photo by Shahram Hesami), e) Alborz (photo by Niloofar Sheik)

### *Loureedia venatica* Henriques sp. n

#### Type material

IRAN, *Alborz Province:* Karaj, Chenarak, 1♂Holotype manual collecting, 8 November 2019, (Currently at ZMUT, but to be permanently deposited at MHNG), A. Beigi leg.; *Alborz Province:* Karaj, Chenarak 1♂ Paratype manual collecting, 8 November 2019, (Currently at SHPC, but to be permanently deposited at MHNG) A. Beigi leg.

#### Additional specimens only recorded by photo

IRAN: *Tehran Province*: 1 ♂, November, 2016, Alireza Zamani (Fig.3.a); Kuhsar, 1 ♂, 11 November 2018, Bayan Golavi (Fig.3.b); 1 ♂, October, 2015, Amir Hossein Bolhari (Fig.3.e); *Kerman Province*: Shahr-e Babak, Meymand Village, 1 ♂, 27 October 2017, Hadis Asghari (Fig.3.c). *Fars Province:* Shiraz, Shahrak-e Sadra, 1 ♂, 2 November 2018, Shahram Hesami (Fig.3.d). *Alborz Province*, Karaj, 1 ♂, 11 November 2016, Niloofar Sheikh (Fig.3.f).

**Fig. 3.**
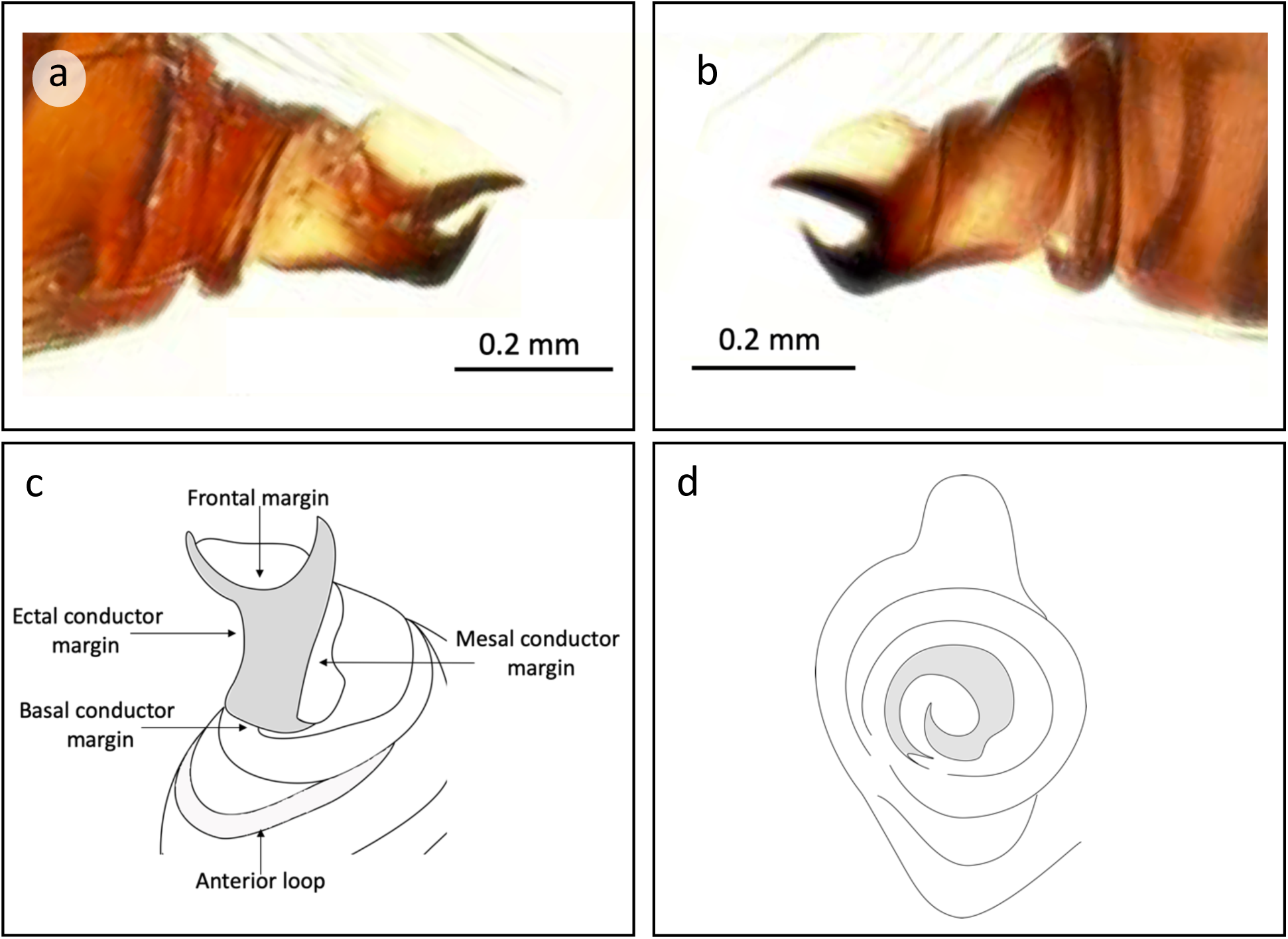
Detailed morphology of the left male pedipalp of *Loureedia venatica* **sp. n**. in a) mesal view, b) ectal view, c) ventral view, d) axial view; a and b, microscope photography of male holotype (photos by Y.Maruzik) c and d schematic illustration of male paratype

#### Etymology

The Latin specific epithet ‘*venatica*’ is the nominative feminine singular of ‘*venaticus*’. Which has been used to describe animals pertaining to hunting. Such as the Asiatic Cheetah (*Acinonyx jubatus venaticus*), a Critically Endangered subspecies which has been studied by a remarkably brave group of Iranian colleagues.

#### Diagnosis

The new species differs from congeners by the dorsal abdominal pattern n and shape of the male palp. The dorsal abdominal pattern is similar to those in *L. annulipes, L. lucasi* and *L. jerbae*, but is distinct by having a compact longitudinal red stripe limited to the medial region (*vs*. dorsal medial region anteriorly black with thin red margins and posteriorly white in *L. annulipes*, and the red stripe extending into lateral projections in *L. lucasi* and *L. jerbae* **comb. n**.). These lateral projections have small compact white dots at their tip in *L. lucasi*, small scattered white dots in *L. jerbae* **comb. n**. or are represented by large white spots in *L. annulipes, vs*. white spots of intermediate size forming lateral projections in the new species. Furthermore, the most anterior pair of white spots is separated in most *Loureedia* species but appears to be often fused or extend very closely to each other in *L. venatica* **sp. n**. and *L. colleni*, occasionally merging with a distinct white spot above the pedicel, forming an anterior white shield (Fig.2 b, c, f), which can be somewhat similar to the anterior coloration in some *L. colleni* males (Henriques, et. al. 2018).

The male pattern has a carapace, sternum, labium, chelicerae, and maxillae dark brown with tones of red. Carapace mostly covered by long black setae and scattered short white setae, also presenting localized patches of short red setae mostly in pars thoracica or the center of pars cephalica. Legs covered with thin black hairs, presenting distinct regions of white hairs at the hedges of all segments, that rarely, or scarcely, connect with each other forming distinct white rings. Abdomen with a compact longitudinal median red stripe with lateral projections having compact white spots at their tip. The most anterior pair of white spots either fused or situated very close to each other, sometimes merging with a distinct white spot above the pedicel, forming an anterior white shield.

When viewed ventrally, the male palp of *L. venatica* **sp. n** shows four specific traits:

1. The mesal margin is straight and slightly bulging outwards; whereas it is highly concave in *L.colleni* and *L.jerbae*, mildly concave in *L.lucasi* and straight in *L.annulipes*.
2. The ectal margin only curves inwards slightly and in a non-smooth way, but rather revealing a flat surface; similarly to *L. colleni* and *L.jerbae* whereas it is highly concave in *L.annulipes* and straight in *L.lucasi*.
3. The dorsal conductor tip is not thicker at its base and is considerably curved, the frontal margin is almost semi-circular causing the ventral tooth to be crescent moon-shaped, facing upwards and reaching outwards almost far as the ventral tooth; whereas it is thicker at the base in *L.lucasi* and *L.annulipes* and straighter i*n L.annulipes* and *L. colleni*.
4. The anterior loop is much longer than in any recorded species, which, on a ventral view, terminates beyond the edge of the conductor’s basal margin. Similarly to *L.jerbae* which can be distinguished by its smaller loop, somewhat similarly to E. lucasi which also as a shorter loop but with a less circular shape, whereas it is clearly much shorter in *L.colleni* and *L.annulipes*.

The male pedipalp of *L.venatica* n.sp. has a very tall basal lamella (when observed in an ectal or mesal view), the dorsal conductor tooth is very smooth and is set at an almost 90° angle from its base, the ventral tooth has a very wide base and is heavily torqued, laterally being perceived as an abrupt 90° bend facing upwards.

Mesal conductor margin almost straight, forming a 90° angle with the basal conductor margin, producing a very small “L” shape. Ectal margin slightly curved inwards. Dorsal conductor tip not thicker at its base and highly curved with a very sharp tip.

The mesal conductor tooth elevation is more easily perceived in an axial view, where it can be noted that it has considerable torsion, which is perceived as very coiled spiral completing its cycle in the direction of the paracymbium tip.

Female. Unknown.

#### Phenology

All available records were made in late October (n=2) and early November (n=5), which matches the known phenology of most other *Loureedia* species (Henriques et. al. 2018). All available records of this species belong to wandering males, but the nest are assumed to be similar in structure to the ones found in other members of this genus (Miller et al. 2012; Henriques et. al. 2018).

#### Habitat

75% records were made at the southern foothills of the Middle Elburz Mountains (n=6) while the remaining 25% of records (n=2) were made on the northern foothills of the East Zagros Mountains (Fig.5), which are overall both semiarid limestone mountains.

**Fig. 4.**
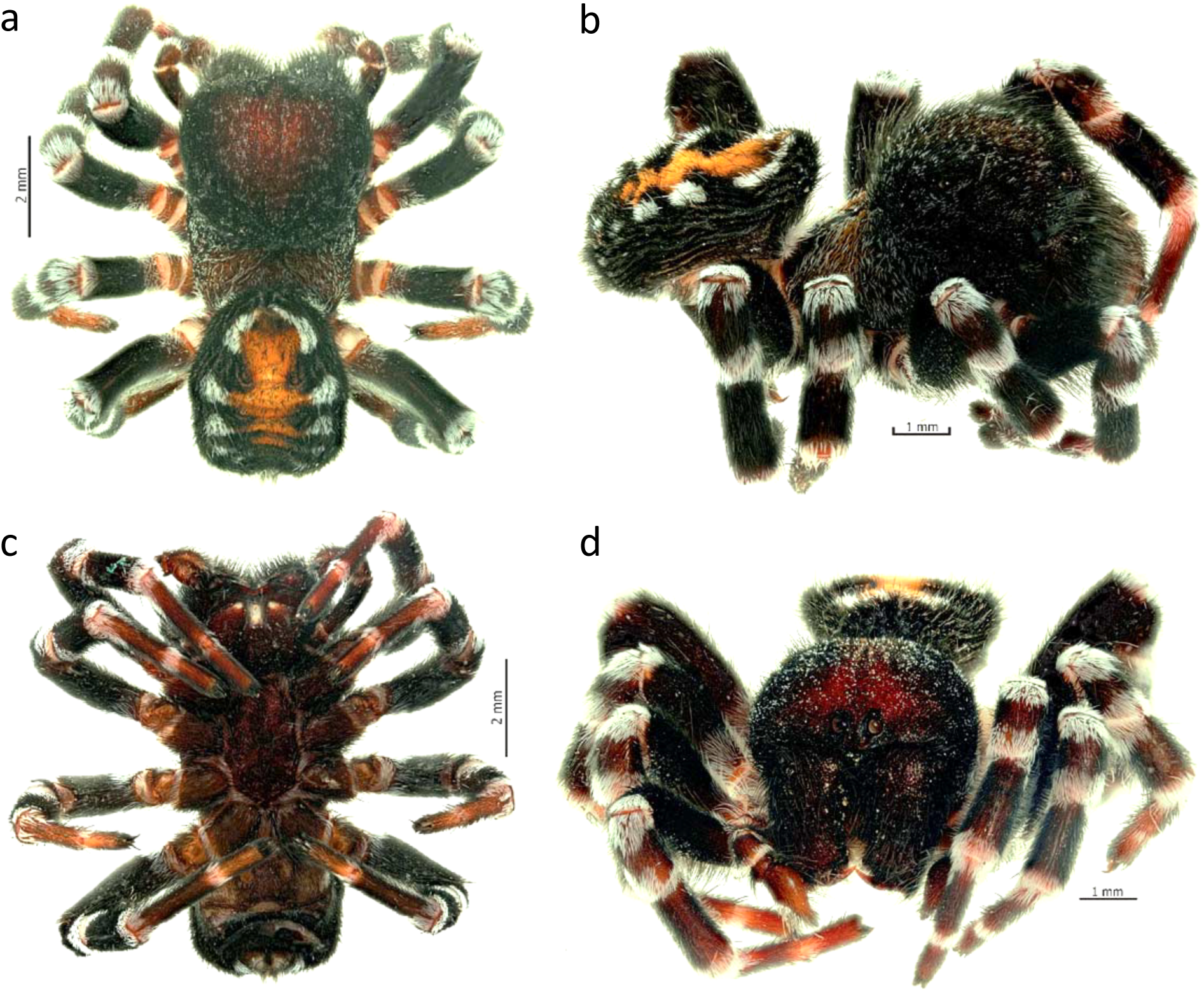
Overall morphology of *Loureedia venatica* n.sp., male holotype in a) dorsal view, b) lateral view, c) ventral view. d) frontal view

**Fig. 5.**
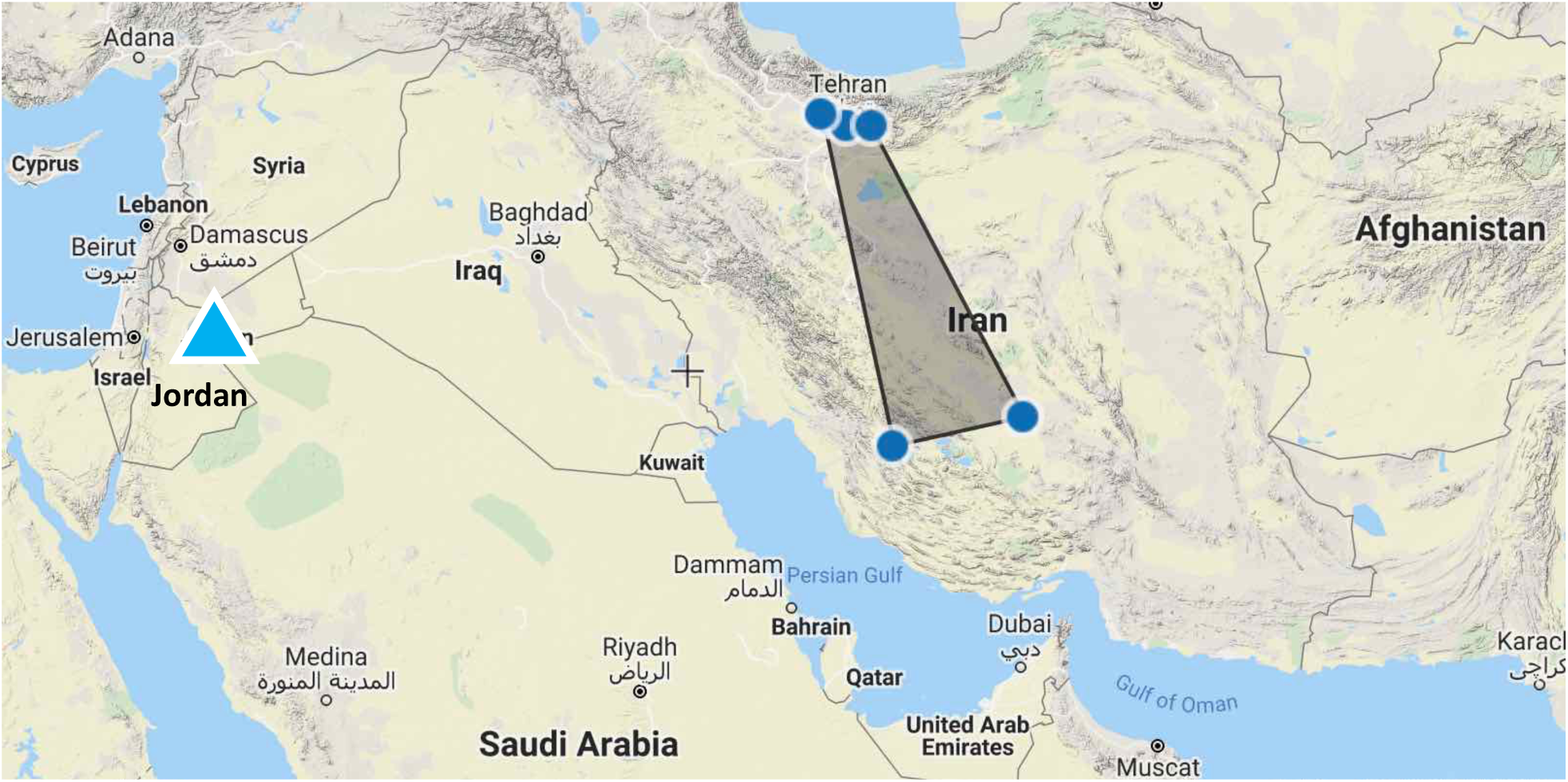
Distribution map of the most Eastwards records of the *Loureedia* genus. With the first record of the genus to Jordan (in light blue triangle) and records of *L.venatica* ***sp. n*. (in** dark blue circles) with the minimum convex polygon of the species range represented by a shaded area (image based on geocat output).

A few of the observations were made in rural or peri-urban areas and around human dwellings, which are unlikely to be the species natural habitat, but are more likely a result of the wandering behaviour of males.

#### Distribution

The Elburz and Zagros mountains are some of Iran’s natural borders and have long been known to have played an important role in shaping the past and present distribution of various Iranian species, limiting ranges between them but also isolating species within them (Macey et al. 2000, Esmaeili-Rineh et al. 2016). Although, from *in vivum* photographic records alone, these Northern (Elburz) and Southern (Zagros) *Loureedia* records appear to be morphologically identical (Fig.3), we cannot exclude the hypothesis that these southern records, which we tentatively place as part of the range of *L.venatica* ***sp. n***., in fact represent a distinct species, as solely northern specimens were analysed in this publication (see type material).

Even under the most encompassing analysis (including all *Loureedia* records from Iran), the Area of Occupancy (AOO) is 24.000 km2 and the Extent of Occurrence (EOO) is 131,646.571 km2 (Fig.5). The impacts of these results and the EOO and AOO values resulting from solely considering northern records, from the point of view of a Red List assessment, are analysed further in the discussion section.

### *Loureedia jerbae* (El-Hennawy, 2005) comb. n., stat. resurr

*Eresus jerbae* El-Hennawy, 2005: 88, figs 1–2 (♀, synonymized with *L. annulipes* by Miller *et al*. [2012]; rejected here).

#### Type material

Holotype, TUNISIA: Djerba, 1♀, [no date], (Latreille), original code 12470, bottle no. 471, tube no. AR 835 (MNHM) originally misidentified as *Eresus petagnae* Audouin, 1826 (currently considered *nomen dubium*).

#### Additional specimens

TUNISIA: Djerba, 1♂manual collecting, 31^st^ October 2019, Macík, S. leg. (LACCUP); Djerba 1 ♀, 2j [no date], (Latreille.), original code 12470, bottle no. 471, tube no. AR 835 (MNHM) originally misidentified as *Eresus petagnae* Audouin, 1826 (currently considered *nomen dubium*).

See comments sections for the exclusion of Algerian records

#### Distribution

Solely know from the Island of Djerba, Tunisia. Likely endemic to that small region.

#### Diagnosis

This species can be differentiated by the characters of the male palp, while its dorsal pattern although different is very similar to *L.lucasi* and other yet undescribed species of North Africa.

When viewed ventrally, the male pedipalp of *L. jerbae* **comb. n** has Trait 1) The mesal margin is irregular, slightly bulging outwards. Trait 2) The ectal margin curves inwards only slightly mostly at the base of the dorsal tooth. Trait 3) The dorsal conductor tip is much thicker at base and is considerably curved. The frontal margin is smooth but curves abruptly close to the tip of the ventral tooth, and the distance between the mesal and the ventral tooth base is very wide, as the distance between the frontal and mesal conductor margins are the widest of any know species. Trait 4) The anterior loop being fairly long and thin, which, terminates beyond the edge of the conductor’s basal margin, but is positioned much closer to it (Fig.7 c).

The male pedipalp has a very thin basal lamella (when observed in a ectal or mesal view), the dorsal conductor tooth is very straight with a triangular shape, with a wide base that acuminates smoothly towards the tip, the ventral tooth faces upwards in smooth gradual angle (Fig. 7 a, b).

**Fig. 6.**
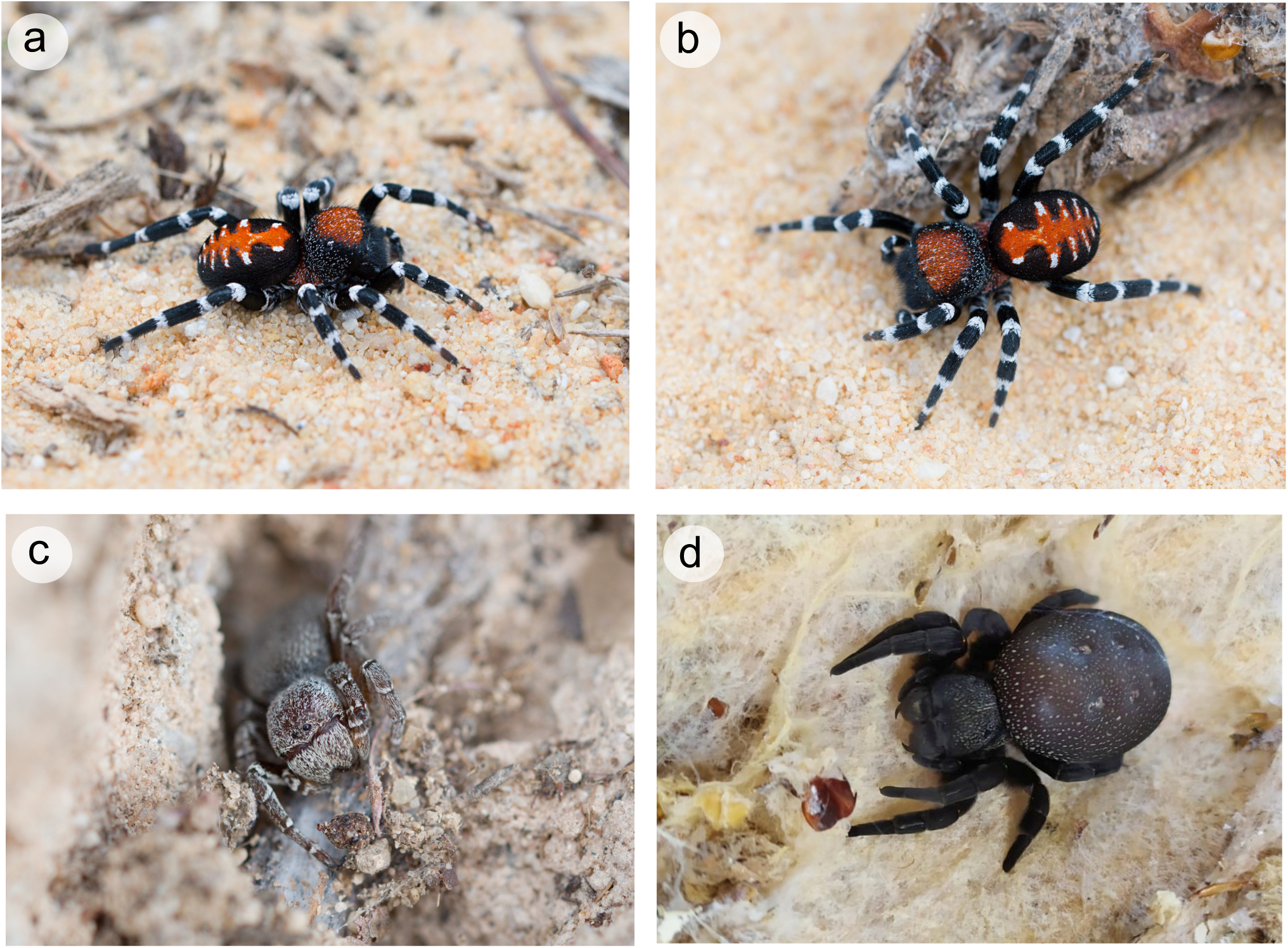
Photographic records of *Loureedia jerbae* **comb. n**. *in vivum*, recording an adult male in a) lateral view and b) dorsal view, as well as c) a juvenile and d) and adult female (photos by Stanislav Macík)

**Fig. 7.**
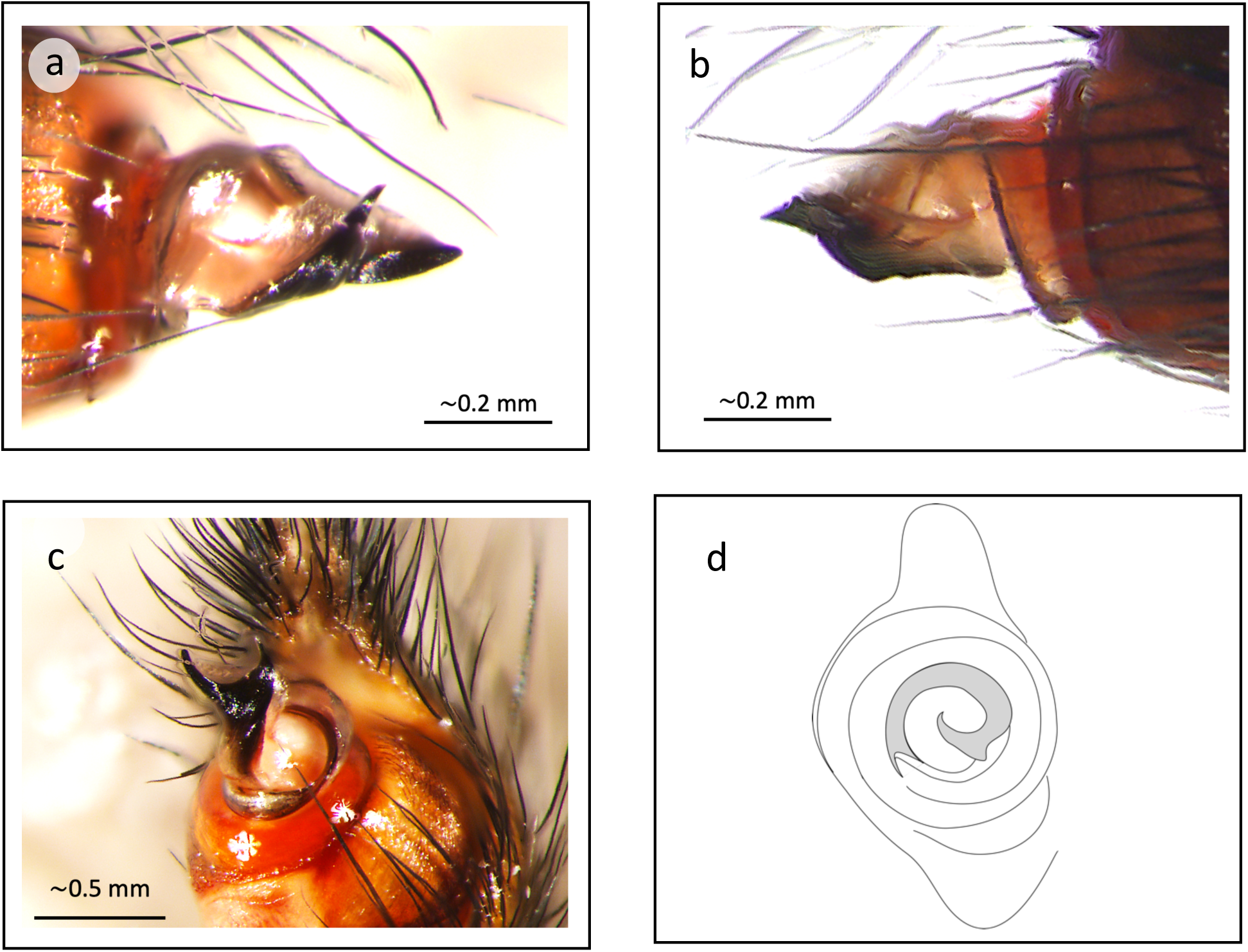
Detailed morphology of male pedipalp of *Loureedia jerbae* **comb. n**. in a) mesal view, b) ectal view, c) ventral view, d) axial view; a, b and c, microscope photography (photos by M.Forman), d schematic illustration of same specimen

The mesal conductor tooth elevation is more easily perceived in an axial view, where it can be noted that it is high torsion, which is perceived as a very tight coiled spiral very close to cymbium centre (Fig. 7 d).

The dorsal abdominal pattern of *L. jerbae* **comb. n** (Fig.6) is similar to the brightly coloured red dorsal patterns present in males of *L. annulipes, L. lucasi* and *L. venatica* **sp. n**., but it appears distinct by having a broad longitudinal red stripe extending into lateral projections (see diagnosis of *L. venatica* **sp**.**n**. for further details). The tip of these lateral projections have small or elongated white dots which can be patchy (Fig. 6 a,b) but can also be thin and somewhat compact (specimens not recorded here). The most anterior white spots are well separated, and the longitudinal red stripe is this area, and the extensions so wide at their base, that it forms a diamond (Fig.6 b).

For the description of the female see El-Hennawy (2005).

#### Comments

The fine structure of the female holotype from Tunisia and the Algerian female (El-Hennawy, 2005) are as distinct from each other than the differences observed between other species of this genus (Henriques *et al*. 2018). Therefore we have not included Algeria in the distribution range of *L. jerbae* **comb. n** which we believe to only be found in the island of Djerba.

#### Pet trade in Loureedia spp

Although the *Loureedia* genus was only recently described (Miller at al. 2012) and so little was known about *L. jerbae* **comb. n**., this species was found to be for sale online, with clear indication of its narrow natural range (Fig.8). We could not ascertain with certainty if *L. venatica* **sp**.**n**. is currently being sold, but photos of this species are currently being used to advertise the sale of *Loureedia* spiders (Fig.9).

**Fig. 8.**
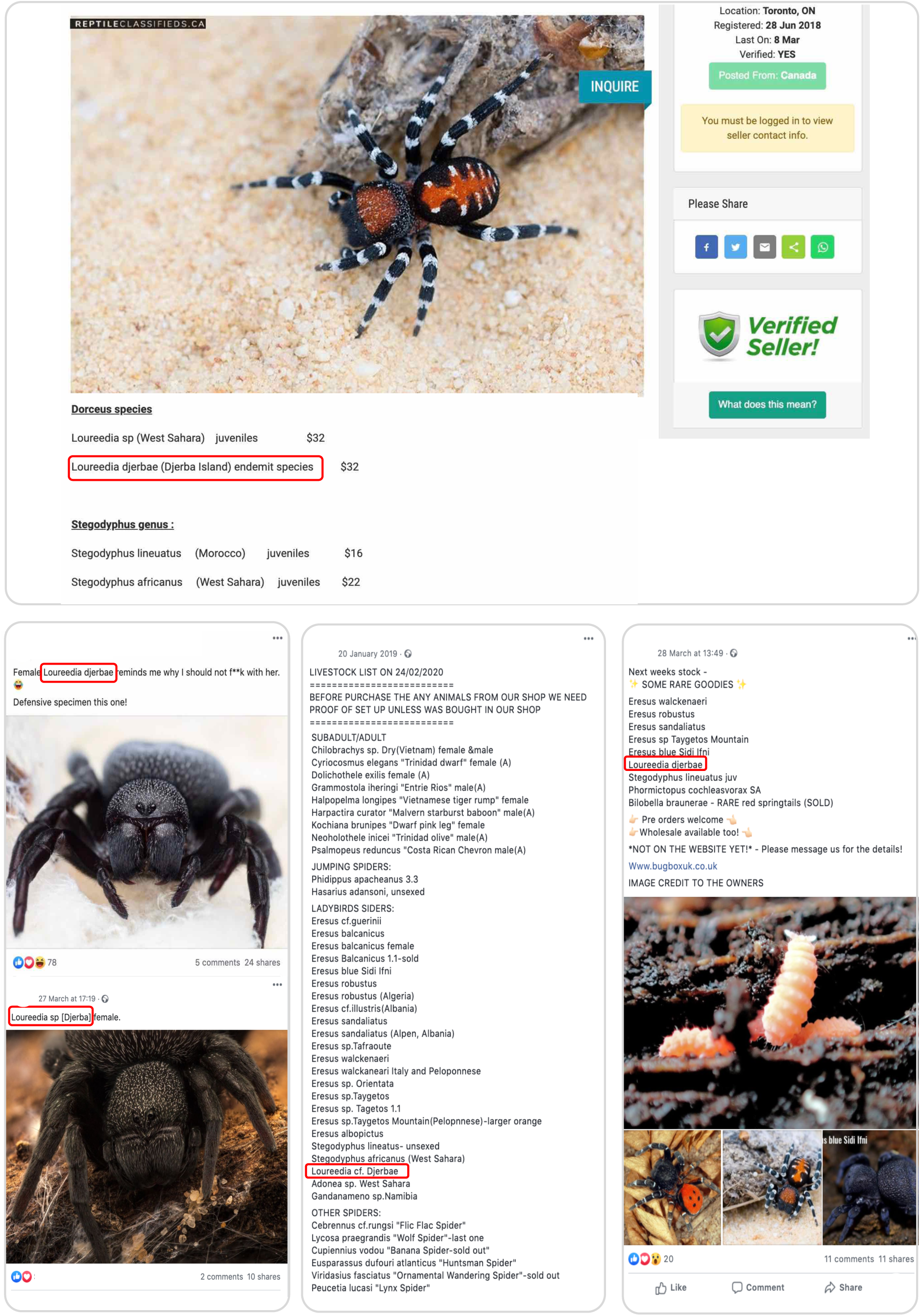
Records of *Loureedia jerbae* ***comb***.***nov***. found in the international pet trade during the first trimester of 2020 alone. All images cropped to hide the identity of those involved, with the references to the species highlighted by a red box. Canadian Website advertising “*Loureedia djerbae*” as an “endemit” species (above), social media posts of three different users based in the UK (bellow).

**Fig. 9.**
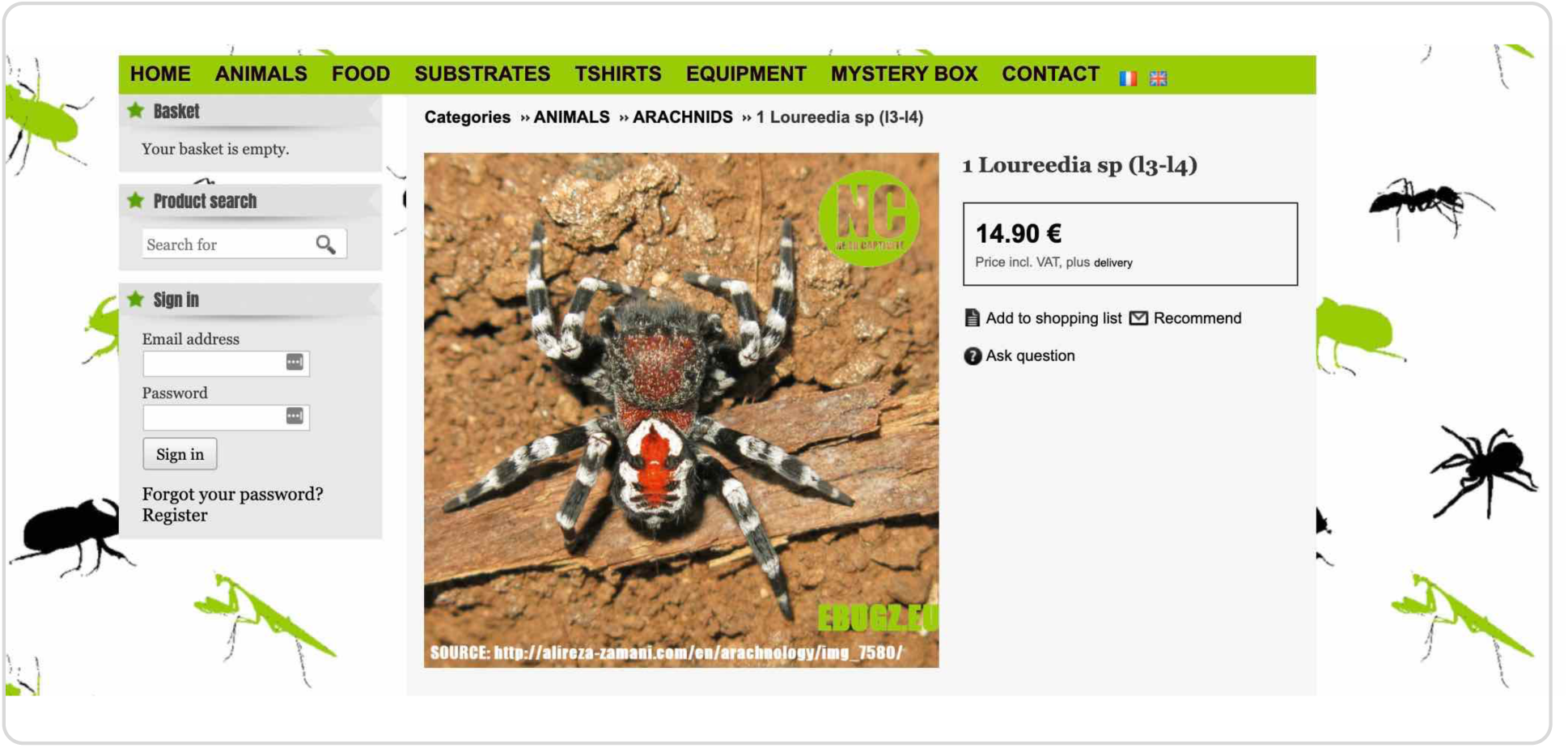
*Loureedia venatica* **sp. n**. used to advertise the sale of its congenerics the international pet trade during the first trimester of 2020 alone. All images cropped to hide the identity of those involved, with the references to the species highlighted by a red box. Canadian Website advertising “*Loureedia djerbae*” as an “endemit” species (above), social media posts of three different users based in the UK (bellow).

In the first trimester of 2020, we found evidence that *Loureedia* species were on sale in France, Germany, Czech Republic and the UK. *L. jerbae* **comb. n**. was also found to be advertised by a Canadian based company (Fig.8), but it is uncertain if the animals are being imported into Canadian territory, or if they were sent from Tunisia (via the EU) directly to US buyers.

## Discussion

The *Loureedia* genus is the only Eresid genera to be described in the twenty-first century, erected from a species previously placed in the *Eresus* genus (Miller at al. 2012). Therefore poorly known *Eresus* species might in fact prove to belong to *Loureedia*, as was the case of *L. lucasi* (Henriques et al. 2018).

Once our analysis focused on Iranian *Loureedia* we therefore considered it essential to revise the taxonomy of *Eresus* species known to that broader region. Which included *E. walckenaeri moerens* C. L. Koch, 1846, recorded from neighbouring Afghanistan (Roewer, 1962) and *E. rotundiceps*, described from Ukraine and reported from Turkmenistan (WSC, 2020). We concluded that these taxa belongs to *Eresus sensu stricto*, and are therefore substantially different from the *Loureedia* genus (Miller at al. 2012) and cannot contain the undescribed female of *L. venatica* **sp. n**.. The in depth analysis of the *Eresus* genus in the region is beyond the scope of this publication and is intended be the focus of an upcoming manuscript in this series.

We found that the species that more closely resembles *L.venatica* **sp**.**n**. is *Eresus jerbae* originally recognized from Tunisia and Algeria. Which is another indication of how easy it can be to generate new synonyms within the *Loureedia* genus, particularly when working eresids, if failing to recognize the historic taxonomic link between these two genus. We resurrect *Eresus jerbae* from its current synonym with *Loureedia annulipes*, while recognizing that it does belong in the same genus, *L. jerbae* **comb. n**. and we describe the male of the species for the first time. Which allowed us to compare and distinguish this species from *L.venatica* **sp**.**n**. (solely known from male specimens). The Algerian record of *Eresus jerbae* (El-Hennawy, 2005) was purposely excluded of our analysis of this species as that specimen contains very distinct features from the holotype of *Loureedia jerbae* **comb. n**., although it is clearly a member of the *Loureedia* genus.

We also looked at the threats and extinction risk metrics of these species. We found that when calculating the range of *L.venatica* **sp**.**n**. based on all available records, this species AOO is very small, and if southern (Zagros) records prove to belong to a different species (see Distribution in the Results section for more details) a reduction analysis shows that there would be a reduction in EOO of almost 100% (from 131,646.571 km2 to 630.843 km2) and a 34% reduction in AOO (from 24.000 km2 to 16.000 km2). Which could potentially place the species distribution within the Endangered category threshold range (following IUCN Red List Criterion B). From a conservation standpoint it is therefore imperative to validate the species identity in its southern (Zagros) range, in order to accurately assess the species extinction risk.

We hope that by contributed towards addressing the Linnaean and Wallacean shortfall in these poorly known taxa, we also highlighted how important those two are to accurately measuring extinction risk. Providing yet another case study of how valuable of an ally taxonomy can be for conservation.

During our research we also found this genus to be present in the pet trade (Fig. 8 and 9). Illegal wildlife trade in invertebrates is rarely a priority of law enforcement, although it often targets high-value rare or endangered species. A trade which unbeknown to many often included spiders (Fukushima et al. 2019).

The retailers importing or selling these animals appeared to be mostly based in Europe and North America. Namely France, Germany, Czech Republic, UK and Canada, which would rarely rank on the top illegal wildlife trade concerns. As fewer animal trade issues are expected in Canada, because although the country is very large, it represents a small portion of the pet trade market, but also because it is illegal to import illegally collected specimens following Canadian law (WAPPRIITA, 1991) which states that: “No person shall import into Canada any animal or plant that was taken, or any animal or plant, or any part or derivative of an animal or plant, that was possessed, distributed or transported in contravention of any law of any foreign state.”

However, although Canadian companies are importing North African species by air cargo via Europe, likely to buyers in the US (Fig. 8). It is impossible to ascertain at this point if these species were actually imported into Canada, or shipped directly from European sellers to the US. The most likely end market of this trade route. Furthermore, in order for Canadian authorities to enforce WAPPRIITA, they need to have evidence that the species were illegally collected. Which although is self-evident by the country of origin of this species, is near impossible to prove, since a species can become “legal stock”, if an egg sac, a pregnant female or even a mating pair, are collected in the wild and the offspring are born in Europe.

From our analysis the online retail of *Loureedia jerbae* from Tunisia to Canada and Europe, and the trade in this genus overall (Fig. 9) appears to be quite prevalent at this point in time, but this might be a bias of our selected snap shot. Where the species was indeed imported from Africa recently, made its way to European and North American markets, but if no further collecting events take place, or the ones planed are stopped by exposing this situation, it might be possible that the *Loureedia* trade will simply die off when the specimens now available die as well. Once that this is what has been observed for other species, when no provision were made to maintain a healthy captive breeding population, or when attempts were made but failed, or become economically unsustainable.

These are just some of the reasons why we strongly propose the long term monitoring of the trade in these small animals to be supported, further research to be conducted, but more importantly for law enforcement to be given more resources to investigate, present substantial cases and eventually even prosecutions. Which includes ascertaining criminal links and compiling historical activities, that we hope this manuscript might have helped with.

## Acknowledgments

Sérgio Henriques is grateful to Robin Freeman, Monika Böhm and Pedro Cardoso for their supervision, as well as Olivia Canteiro-Henriques and Cátia Canteiro for her unconditional support.

We are sincerely grateful to Dr. Luca Bartolozzi (curator, UNIFI), for arranging the loan of eresid specimens under his care. To the Muséum National d’Histoire Naturelle, Paris and in particular to Dr. Piotr Daszkiewicz from for assistance with historical information, translations and advice, to Dr. Christine Rollard and Christophe Hervé for support and access to the arachnological collection. To the Museum & Institute of Zoology PAS, and in particular to Wioletta Wawer for assistance with historical information, translations and access to arachnological collection. To Janet Beccaloni, Duncan Sivell and Daniel Whitmore for their friendship and help assessing correspondence, records and specimens held at the Natural History Museum, London. To Maria Mostadius and Jonas Ekström at the Biologiska Museet Lund for their assistance in analysing the specimens from the Lindberg collection. To Nina Polchaninova, for her help and support of the analysis of this family in the region. To Jason Dunlop, Curator of arachnids and myriapods at the Museum für Naturkunde, Leibniz Institute for Evolution and Biodiversity Science, Berlin for information on Koch’s collection and always useful and lively discussions. To Peter Jaeger and Julia Altman for their continued support of my work on this family, and access to the specimens held at the Senckenberg Museum. To Martin Forman of the Laboratory of Arachnid Cytogenetics at Charles University in Prague, Milan Řezáč of Crop Research Institute in Prague and in particular to Stanislav Macík, for their support of the revision of this group in the region.

To Amir Beigi for providing with the collected material; Majd al Qadi, Hadis Asgari, Amir Hossein Bolhari, Bayan Golavi, Shahram Hesami, Mohammad R. Shakhatreh and Niloofar Sheik for providing us with their photographic records; and Atefe Shayanmehr and Reza Vasighi for field assistance.

We are also grateful to Ernest Cooper and Rick West for their help and input towards the analysis of this species presence in the pet trade.

